# Leukemia inhibitory factor (LIF) receptor amplifies pathogenic activation of fibroblasts in lung fibrosis

**DOI:** 10.1101/2024.05.21.595153

**Authors:** Hung N. Nguyen, Yunju Jeong, Yunhye Kim, Yaunghyun H. Kim, Humra Athar, Peter J. Castaldi, Craig P. Hersh, Robert F. Padera, Lynette M. Sholl, Marina Vivero, Nirmal S. Sharma, Jeong Yun, Louis T. Merriam, Ke Yuan, Edy Y. Kim, Michael B. Brenner

## Abstract

Fibrosis drives end-organ damage in many diseases. However, clinical trials targeting individual upstream activators of fibroblasts, such as TGFβ, have largely failed. Here, we target the leukemia inhibitory factor receptor (LIFR) as a “master amplifier” of multiple upstream activators of lung fibroblasts. In idiopathic pulmonary fibrosis (IPF), the most common fibrotic lung disease, we found that lung myofibroblasts had high LIF expression. Further, TGFβ1, one of the key drivers of fibrosis, upregulated LIF expression in IPF fibroblasts. In vitro anti-LIFR antibody blocking on human IPF lung fibroblasts reduced induction of profibrotic genes downstream of TGFβ1, IL-4 and IL-13. Further, siRNA silencing of LIFR in IPF precision cut lung slices reduced expression of fibrotic proteins. Together, we find that LIFR drives an autocrine positive feedback loop that amplifies and sustains pathogenic activation of IPF fibroblasts downstream of multiple external stimuli, implicating LIFR as a therapeutic target in fibrosis.

**Significance Statement:** Fibroblasts have a central role in the pathogenesis of fibrotic diseases. However, due to in part to multiple profibrotic stimuli, targeting a single activator of fibroblasts, like TGFβ, has not yielded successful clinical treatments. We hypothesized that a more effective therapeutic strategy is identifying a downstream “master amplifier” of a range of upstream profibrotic stimuli. This study identifies the leukemia inhibitory factor receptor (LIFR) on fibrotic lung fibroblasts amplifies multiple profibrotic stimuli, such as IL-13 and TGFβ. Blocking LIFR reduced fibrosis in ex vivo lung tissue from patients with idiopathic pulmonary fibrosis (IPF). LIFR, acting as a master amplifier downstream of fibroblast activation, offers an alternative therapeutic strategy for fibrotic diseases.

## Introduction

Fibroblasts are the main producers and drivers of collagen production, the largest component of the extracellular matrix and primary contributor to fibrosis. TGFβ (1), IL-4, and IL-13 (2) are major stimulators of collagen production by fibroblasts. TGFB1, one of three isoforms in the TGFβ family with pleiotropic functions across areas of development, cell proliferation, and regulation of inflammation, plays a prominent role in fibrosis (1, 3). In IPF and other fibrotic lung diseases, myeloid and epithelial cell subpopulations are major sources of secreted TGFβ1 (4). TGFβ1 signaling proceeds via several pathways, both canonical and non-canonical (5). In the canonical pathway, binding of TGFβ1 to its receptor phosphorylates the SMAD2 and SMAD3 transcription factors, which then drive profibrotic transcriptional programs in collagen synthesis and myofibroblast differentiation. Furthermore, TGFβ1 is able to amplify its profibrotic effect by inducing the gene expression of other drivers of fibrosis, such as IL-11 (6). TGFβ signaling in fibroblasts can be further modified via a number of different mechanisms, including other cytokines (such as trans signaling via IL-6 (7)), Hippo and other signaling pathways (8), metabolic shifts (9), non-coding RNAs (10), and others (3). Pirfenidone, one of two clinically approved medications for IPF, is thought to act (in part) by inhibiting the TGFβ axis (11). Like TGFβ1, IL-13 is also a prominent upstream inducer of airway (12) and lung fibrosis (13) and is a candidate therapeutic target in lung fibrosis (14). IL-13 acts both directly on fibroblasts, as well as on other cell types. For example, IL-13 can promote proliferation of lung fibroblasts via upregulation of autocrine PDGF signaling (15) and drive fibroblast and epithelial cell differentiation towards a myofibroblast phenotype (16). While important in homeostatic contexts (e.g., wound healing (2, 17)), TGFb, IL-4 and IL-13 cytokines play prominent roles across a range of fibrotic diseases, such as fibrotic ILD, autoimmune diseases (e.g., systemic sclerosis), and inflammatory bowel disease (1). However, to date, clinical trials targeting epithelial activators of TGFβ (αvβ6), IL-4, or IL-13 (18–21) in IPF have not shown success.

Interstitial lung disease (ILD) affects up to 0.2% of people and contributes to 0.7% of total deaths (22). While many diseases affect the lung interstitium (i.e., the extracellular and extravascular space), the highest levels of morbidity and mortality are seen in fibrotic ILD. Idiopathic pulmonary fibrosis (IPF) is the most common and well described fibrotic ILD, characterized by progressive lung fibrosis, worsening dyspnea, reduced lung volume, and decreased oxygenation. IPF remains incurable, with a median patient survival of only 3 to 5 years (23). Current anti-fibrotic treatments for IPF only modestly slow the rate of fibrosis and have limited effects on outcomes (24). Accumulating a deeper understanding of the pathobiology of progressive fibrotic ILD is critical for developing more effective pharmacological therapies.

In view of the disappointing clinical results in targeting upstream inducers of fibrosis like TGFβ1 or IL-13, an alternative strategy may involve targeting downstream genes that integrate the effects from multiple upstream stimuli. Such autocrine amplifier pathways may be used in the context of multiple different stimulators, offering an advantage as a therapeutic target. In this study, we asked what mechanisms in lung fibroblasts amplify upstream external, profibrotic stimuli in IPF fibroblasts. We found increased expression of leukemia inhibitory factor (LIF) in the lung fibroblasts of IPF patients, and that in vitro TGFβ1 induced expression of LIF in IPF lung fibroblasts. Given our previous findings that LIFR was an autocrine amplifier of inflammation in fibroblasts (25), we hypothesized that LIFR might mediate an autocrine amplification loop in IPF lung fibroblasts that drives profibrotic gene programs. We tested the effects of in vitro LIFR blockade and gene silencing in primary IPF lung fibroblasts and human IPF precision cut lung slices (PCLS) following stimulation with multiple profibrotic stimuli. We demonstrate that autocrine signaling by the LIF-LIFR axis is a master amplifier of multiple exogenous profibrotic stimuli in IPF lung fibroblasts.

## Results

### LIF is upregulated in lung fibroblasts from IPF patients

We screened for amplifiers of profibrotic pathways in fibroblasts from IPF patients. Ghosh et al. had identified that the gene set for the “IL6 JAK STAT3” pathway has increased expression in IPF lung compared to control (26). In bulk RNA-sequencing (RNA-seq) of lung tissue from IPF patients (n=231) and control (n=267) (26), we examined the expression of genes in the IL-6 family of cytokines. Expression of IL6, OSM, and CTF1 were reduced in IPF lung, while expression of LIF was increased compared to control (Log2 fold change = 0.41, p<0.0001) (Figure 1A). Next, we identified the cell types expressing LIF in IPF lung. In the single-cell (sc)RNA-seq dataset of IPF lung published by Adams et al (27), fibroblasts had the strongest expression of LIF compared to epithelium, endothelium, and immune cells (Figure 1B). Furthermore, lung myofibroblasts from IPF lungs had increased LIF expression compared to myofibroblasts from control lung in single-cell analysis (Figure 1C). In early passage, primary lung fibroblasts from IPF patients, we found that stimulation with TGFβ in vitro increased LIF expression by over 5-fold (p<0.01, Figure 1D). Taken together, these findings suggested that LIF may play a pathogenic and profibrotic role in IPF lung fibroblasts.

**Fig. 1.**
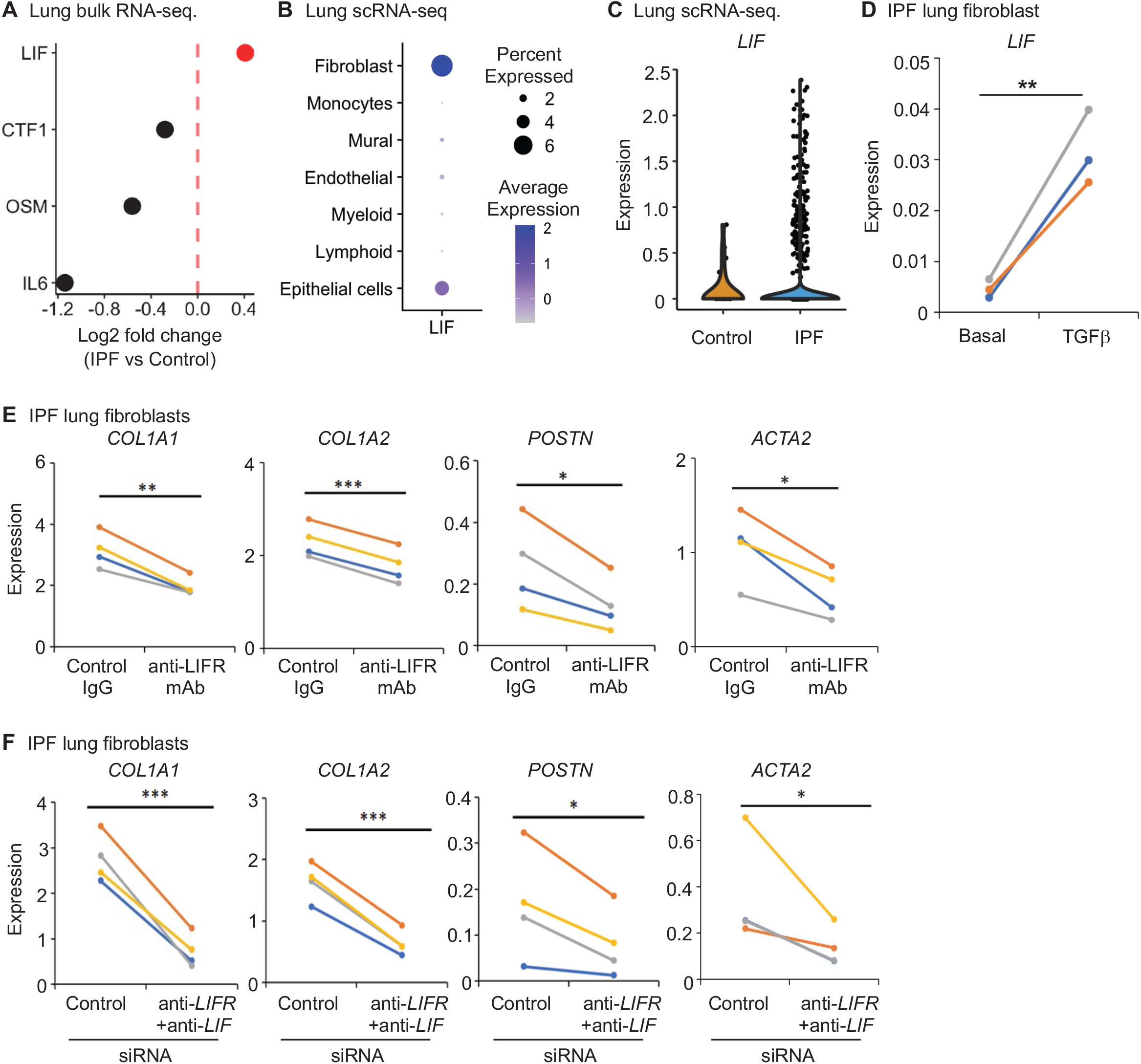
*LIF* is upregulated in lung fibroblasts from IPF patients, and LIFR drives the induction of profibrotic genes by TGFβ. (A) Gene expression in bulk RNA-seq of lung tissue from IPF patients (n=231) compared to control (n=267). Genes in the IL6 family with significantly different expression between IPF and healthy cohorts are shown. (B-C) Analysis of the single-cell RNA-seq dataset of IPF lung tissue from Adams et al. (27) (B) Dot plot of LIF expression in pseudobulk analysis of cell lineages in human IPF lung. (C) Violin plot of LIF expression in lung myofibroblasts from control or IPF patients. (D-F) Early passage, primary lung fibroblasts were derived from IPF patients. Each line represents a different patient. Data represent gene expression in fibroblasts at 24 h after stimulation by TGFβ1 as measured by qPCR and normalized to GAPDH. (D) IPF lung fibroblasts were incubated with media only (basal) or TGFβ1 (5ng/ml) and LIF expression measured by qPCR (relative to GAPDH). (E) Lung fibroblasts were stimulated with TGFβ1 (5ng/ml) in the presence of an antibody against LIFR (LIFR mAb) or its isotype control (Ctrl IgG). (F) Lung fibroblasts were transfected with siRNAs against LIFR and LIF (LIFR + LIF) or a control (Ctrl) siRNA. 48 h after transfection with siRNA, cells were stimulated with TGFβ1 (5ng/ml). (D-F) Paired Student’s t test, *p < 0.05, **p < 0.01, ***p < 0.001, ns: not significant.

### Blockade of LIFR suppressed profibrotic genes induced by TGFβ1

Given the increased expression of LIF in IPF lung fibroblasts, we hypothesized that LIF and its receptor LIFR on lung fibroblasts can form an autocrine loop amplifying fibrosis. We examined early passage, primary lung fibroblasts from IPF patients in vitro. TGFβ1 was used as an upstream “signal 1” inducing a profibrotic gene program (Supplemental Figure S1). In the presence of the LIFR blocking mAb, the induction of profibrotic genes including COL1A1, COL1A2, POSTN and ACTA2 by TGFβ1 was significantly reduced, compared to an isotype control antibody (Figure 1E). IL-11 is a therapeutic target in IPF that increases expression of profibrotic proteins by fibroblasts and promotes myofibroblast differentiation (28). Like LIF, IL-11 is a member of the IL-6 family and its expression is induced by TGFβ1(6, 28). However, blocking LIFR did not affect the expression of IL11 (Supplemental Figure S2). To confirm the role of a LIF / LIFR autocrine loop, we used siRNAs to silence the expression of both LIF and LIFR in IPF lung fibroblasts (Supplemental Figure S3). Fibroblasts transfected with LIF and LIFR siRNA had reduced expression of TGFβ1-induced profibrotic genes compared to cells transfected with control siRNA (Figure 1F), with similar observations with a different LIFR siRNA (Supplemental Figure S3A). Several other profibrotic genes induced by TGFβ1 were also significantly suppressed by silencing of LIF and LIFR, such as COL3A1, COL5A2, COL6A2, COL6A3, MMP2, CTGF, and FN1 (Supplemental Figure S4). Taken together, these results support that LIF and its receptor LIFR form an autocrine loop in fibroblasts that amplifies the induction of profibrotic genes by the upstream stimulus TGFβ1.

### LIFR signaling is required by major profibrotic activators

We next asked if the autocrine LIF/LIFR loop is also required by other upstream, profibrotic activators. We used siRNA to silence the expression of both LIF and LIFR in IPF lung fibroblasts and compared to transfection with control siRNA. Silencing LIF and LIFR significantly reduced the induction of collagen 1A1, periostin, and α-smooth muscle actin protein by IL-4, IL-13, or TGFβ1 (Figure 2), with similar results from a different LIFR siRNA (Supplemental Figure S5). These results support that LIF and LIFR form an autocrine loop in fibroblasts that is a master amplifier of many profibrotic stimuli.

**Fig. 2.**
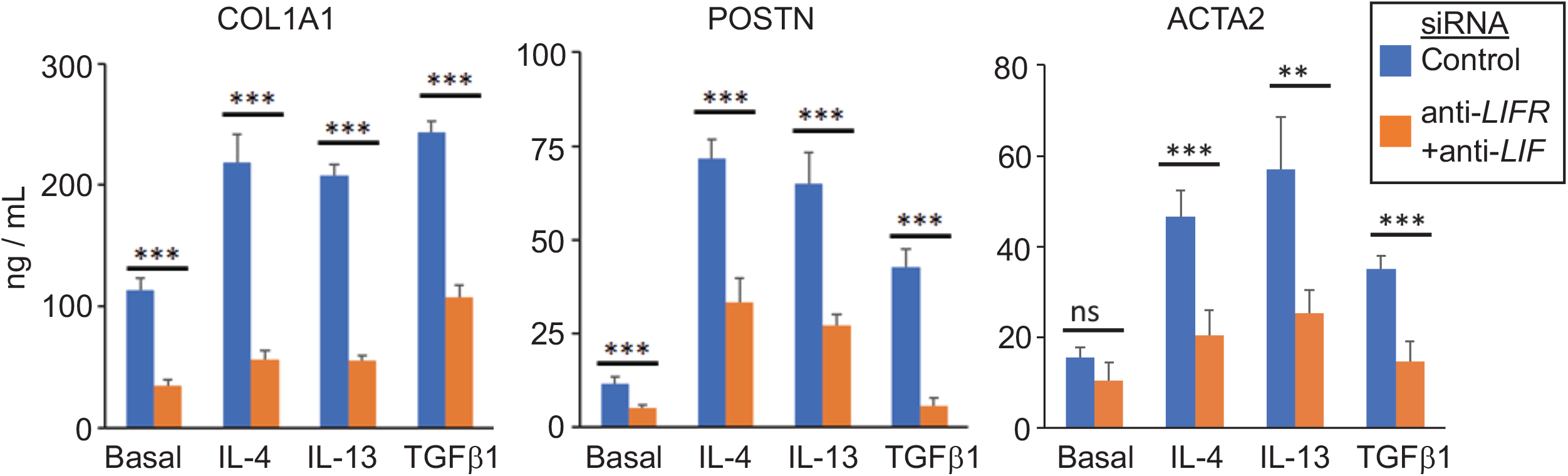
LIFR signaling by lung fibroblasts amplifies profibrotic genes induced by IL-4, IL-13 or TGFβ1. Primary IPF lung fibroblasts were transfected with siRNAs against LIFR and LIF (LIFR + LIF) or a control (Ctrl) siRNA. 48 h after transfection with siRNA, cells were stimulated with IL-4 (50ng/mL), IL-13 (50ng/mL), TGFβ1 (5ng/ml), or left unstimulated (basal). The protein levels were measured by ELISA in cell lysates at 72 h post-stimulation. Error bars represent SD of quadruplicate technical replicates. One-way ANOVA test (**p < 0.01, ***p < 0.001).

### JAK2 mediates the induction of profibrotic genes by TGFβ, IL-4, and IL-13

After binding LIF, LIFR dimerizes with the co-receptor GP130 (IL6ST) to activate a JAK kinase signaling cascade that varies by context. LIF activates JAK1 and TYK2 in synovial fibroblasts (25) but activates JAK1 and JAK2 in cardiomyocytes (29). To examine the mechanism by which LIFR induces profibrotic genes, we utilized the small molecule JAK inhibitors in clinical use for autoimmune diseases and, in the case of baricitinib, COVID-19 pneumonia. We used ruxolitinib and baricitinib, which inhibit JAK1 and JAK2, and tofacitinib, which inhibits JAK3 at lower concentrations, and JAK1 at higher concentrations (30–32). We titrated the doses of these small molecule inhibitors to maximize their JAK selectivity. We tested these inhibitors in vitro on primary IPF lung fibroblasts stimulated with TGFβ1, IL-4 and IL-13. We observed that the inhibitors of JAK1 and JAK2, baricitinib and ruxolitinib, displayed a dose-dependent inhibition of COL1A1 and ACTA2, while tofacitinib did not (Figure 3A).

**Fig. 3.**
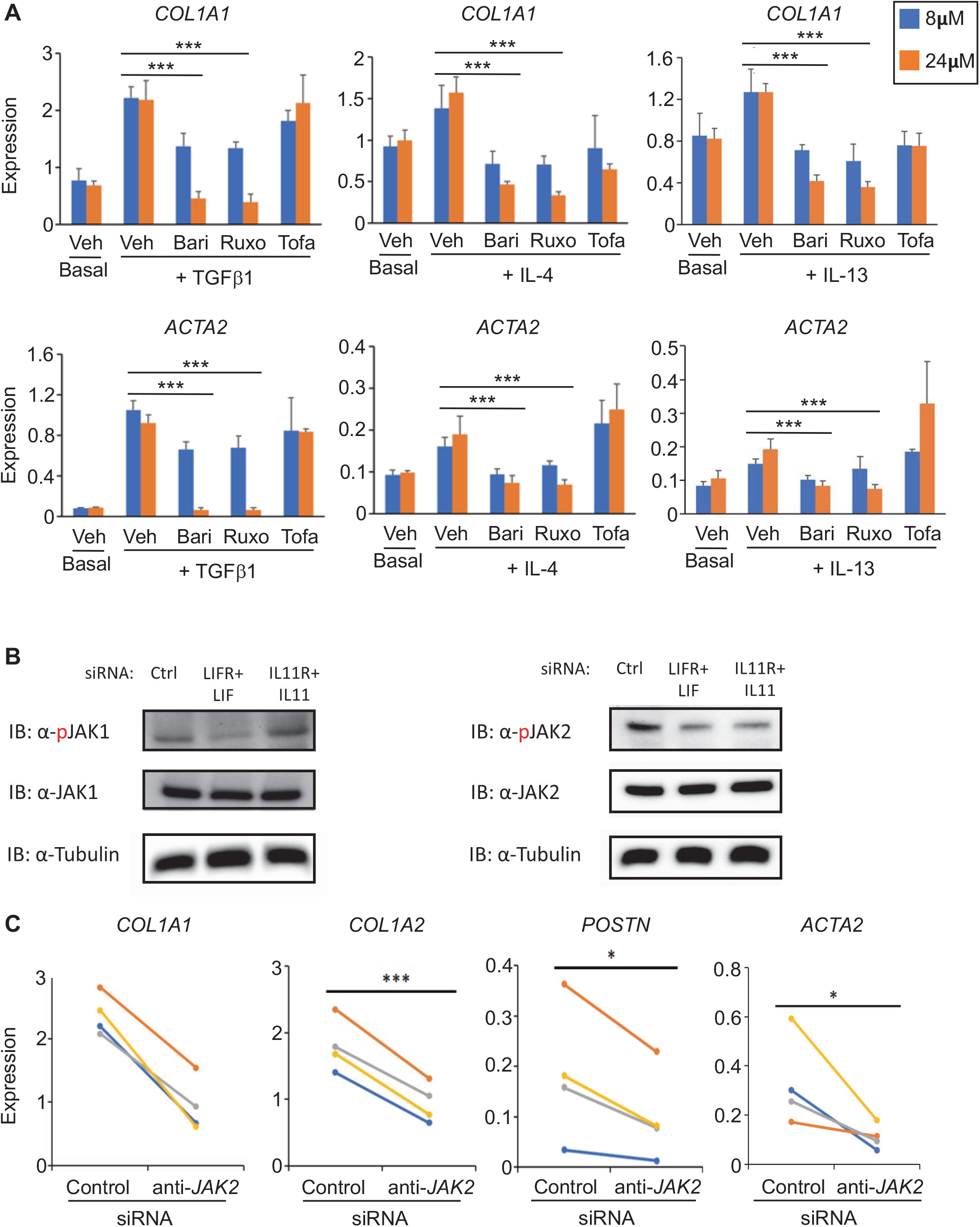
JAK2 mediates the induction of profibrotic genes by TGFβ1, IL-4, and IL-13 in lung fibroblasts. (A) Primary IPF lung fibroblasts were treated with 8 or 24μM of JAK inhibitors or left untreated (basal). Cells were stimulated 0.5 h later with TGFβ1 (5ng/ml), IL-4 (50ng/mL), or IL-13 (50ng/mL). 48 h after stimulation, COL1A1 and ACTA2 expression (normalized to GAPDH) were measured by qPCR. Veh = Vehicle control; Bari = Baricitinib; Ruxo = Ruxolitinib; and Tofa = Tofacitinib. (B) Primary IPF lung fibroblasts were transfected with control siRNA (Ctrl), with siRNA against LIFR and LIF (LIFR + LIF), or with siRNA against IL11R and IL11 (IL11R + IL11). Cell lysates 72 h after siRNA transfection were run on SDS-PAGE gel with immunoblotting with antibodies specific for phosphorylated (p)JAK1 (Tyr1034/1035), JAK1, pJAK2 (Tyr1007/1008), JAK2, and beta-tubulin (loading control). (C) Primary IPF lung fibroblasts were transfected with a control (Ctrl) siRNA or a siRNA against JAK2. 48 h after siRNA transfection, fibroblasts were stimulated with TGFβ1 (5ng/ml), and 24 h later gene expression (normalized to GAPDH) was measured by qPCR. (A) Error bars represent SD of quadruplicate technical replicates. (C) Paired Student’s t test. *p < 0.05, **p < 0.01, ***p < 0.001.

This finding suggested TGFβ1, IL-4, and IL-13 may all share the requirement of either JAK1 or JAK2 for full induction of profibrotic genes. Since TGFβ1, IL-4, and IL-13 also required LIFR for full induction of profibrotic genes (Figure 2), we hypothesized that JAK1 or JAK2 is critical for transducing LIFR signaling. We utilized siRNA against LIFR and LIF or siRNA against IL11R and IL11 to silence LIFR or IL11R signaling, respectively, in primary IPF lung fibroblasts. Silencing of LIFR signaling resulted in a reduction of both phospho-JAK1 (pJAK1) and phospho-JAK2 (pJAK2). In contrast, silencing of IL11R signaling only resulted in a reduction of phospho-JAK2 (pJAK2) (Figure 3B). Thus, JAK2 is a JAK kinase shared downstream of two distinct profibrotic pathways in IPF lung fibroblasts.

Next, we tested the requirement of JAK2 for induction of profibrotic genes. Lung fibroblasts transfected with an siRNA against JAK2 had significantly reduced expression of COL1A1, COL1A2, POSTN, and ACTA2 after stimulation by TGFβ1 (Figure 3C), with similar results from a different siRNA against JAK2 (Supplemental Figure S6). These results demonstrate that JAK2, which is downstream of both LIFR and IL-11R, has a critical role in IPF lung fibroblasts; and targeting JAK2 can interrupt the induction of profibrotic genes.

Blocking LIFR signaling reduces fibrosis in human IPF lung tissue We used primary IPF lung fibroblast lines in vitro to isolate the autocrine functions of LIF and LIFR (Figures 1-3). Next, we tested the role of LIFR signaling in the full cellular context of the IPF lung using precision cut lung slices (PCLS) from IPF patients. PCLS are intact slices of lung tissue explanted from an IPF patient undergoing lung transplantation. PCLS preserves the lung architecture of airways and alveoli in vitro (Figure 4). siRNA against LIFR effectively silenced expression of LIFR in IPF PCLS (Figure 4A). Silencing of LIFR in IPF PCLS resulted in significant reduction of multiple collagen genes including COL1A1, COL1A2, COL3A1 and COL6A3, while not affecting expression of STAT3 (Figure 4A). Knockdown of LIFR also reduced protein expression of COL1A1 and FN1 in IPF PCLS (Figure 4B). These results suggest that LIFR is a key amplifier of the expression of fibrotic genes in IPF lung.

**Fig. 4.**
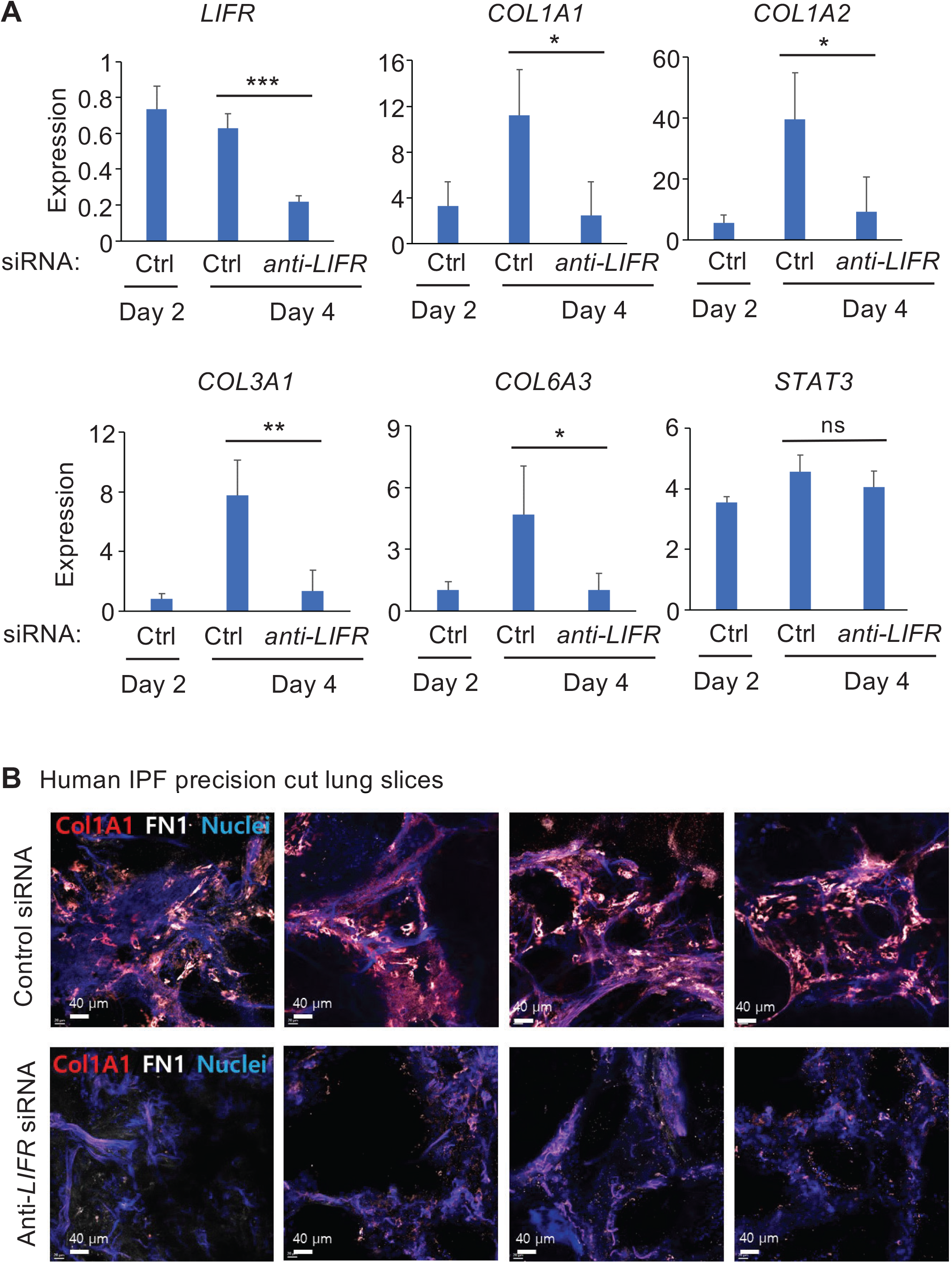
Targeting LIFR in human precision cut lung slices (PCLS) reduces expression of fibrotic genes. Human IPF PCLS were incubated with control (Ctrl) siRNA or siRNA against LIFR on day 0. (A) On days 2 and 4, gene expression (normalized to HPRT) were measured by qPCR. (B) On day 4, PCLS were imaged by immunofluorescence with mAb specific for COL1A1 and FN1. (A) Mean + SD shown from 4 slices per condition. Unpaired Student’s t-test. *p<0.05, **p<0.01, ***p<0.001, ns = non-significant.

## Discussion

In this study, we found that LIF was markedly upregulated in lung myofibroblasts from IPF patients. We demonstrate that LIFR drives an autocrine loop that amplifies the expression and translation of profibrotic genes induced by three prominent upstream stimuli of fibrosis: TGFβ1, IL-4, and IL-13. LIFR acted independent of IL11 expression, but shared downstream signaling via phosphorylation of JAK2. In a clinically relevant approach, we found that in situ siRNA silencing of LIFR in human IPF PCLS markedly reduced expression of collagen and fibronectin. The finding of an LIFR autocrine loop in IPF that amplifies multiple upstream pro-fibrotic stimuli represents a new and attractive therapeutic target that may be effective no matter what upstream drivers of lung fibrosis are actively present.

Literature to date has largely focused on LIF’s paracrine effect in cancer and tumor immunity (33, 34). However, there is a growing appreciation for the pleiotropic nature of LIF and its receptor in other diseases. Some reports describe a protective role for LIF in murine models of organ injury and fibrosis. For example, activation of LIFR on hepatocytes (35) and adipocytes (36) has been shown to ameliorate liver steatosis. After myocardial infarction, LIF protected cardiomyocyte survival and improved cardiac function without inducing fibrosis (37, 38). Yet, other studies support a role for LIF in ameliorating fibrosis since LIF inhibited collagen production in renal fibroblasts, and LIF reduced renal tubulointerstitial fibrosis in a mouse model of unilateral ureteral obstruction (UUO) (39). Others have found that a LIF-neutralizing antibody reduced renal tubulointerstitial fibrosis in experimental UUO, attributed to fibroblast production of LIF that acted on renal proximal tubular cells (40). Hence, a careful examination of the end-organ and disease-context is required to understand the role of LIFR.

Autocrine signaling loops are initiated when a cell secretes a ligand that subsequently binds to and activates its respective receptor on the same cell. Prominent autocrine signaling loops range from EGFR on cancer cells (41) to the IL-2 receptor on T cells (42). Widely described across normal and disease contexts, autocrine loops among cytokines are a consistent feature in the pathobiology of fibrosis. For example, TGFβ has demonstrated autocrine signaling activity in wound healing, experimental models of tissue fibrosis, and tumor fibrogenesis (43, 44). Here, we suggest that the LIF autocrine mechanism involves a sequential two-step process: The initial external upstream “signal 1” can be driven by any one of a range of stimuli (e.g., TGFβ1, IL-4, IL-13) that induces expression of LIF in fibroblasts. Then, in “signal 2,” LIF binds to LIFR on the same fibroblast, creating an autocrine loop that amplifies downstream fibrotic gene programs (Supplemental Figure S7). Targeting the LIFR mediated autocrine loop in fibroblasts can inspire several clinical implications. First, a major challenge in treating IPF is the likely pathophysiology that a range of risk factors produce several upstream stimuli that drive activation of fibroblasts (45). Thus, rather than targeting just one of many upstream stimuli, a more effective therapeutic strategy may be targeting a central downstream amplifier, such as LIFR, that integrates multiple upstream stimuli. Second, autocrine loops can potentially support a “runaway” positive feedback loop that can self-perpetuate, even without continuation of the initial “signal 1”, such as the worsening of IPF that can occur following lung cancer resection (46). Thus, targeting LIFR signaling may be effective in many contexts of fibrosis.

## Materials and Methods

### Cell Lines and Reagents

Human lung tissue was obtained from patients with progressive fibrotic ILD during surgical lung biopsy or lung explantation (N=4). After disaggregation, fibroblasts were isolated using Liberase TL (Sigma), elastase (Worthington Biochemical Corporation), hyaluronidase (Worthington Biochemical Corporation), DNase I (Sigma) as in our previous studies (47, 48). Each unique cell line was derived from a single donor and used experimentally between passages 5 and 8. Cells were maintained in Dulbecco’s modified Eagle’s medium (DMEM) supplemented with 10% fetal bovine serum (FBS; Gemini), 2 mM L-glutamine, 50 µM 2-mercaptoethanol, antibiotics (penicillin and streptomycin), and essential and nonessential amino acids (Life Technologies). Human precision cut lung slices (PCLS) were purchased from IIVS and were grown in DMEM/F-12 media (Gibco) supplemented with antibiotics and antimycotic reagents.

The following antibodies were used: anti-LIFR (clone 1C7), control IgG1 (MOPC21 clone), anti-α-tubulin (Sigma); anti-pJAK1, anti-pJAK2, anti-COL1A1 (Cell Signaling Technologies). Other reagents were purchased from the following vendors: COL1A1 (Collagen 1A1), POSTN (Periostin) ELISA kits (R&D Systems); ACTA2 (aSMA) ELISA kit (Abcam); TGFb, IL-4, IL-13 (Peprotech). siRNAs were purchased from Life Technologies (for human cell lines) and from Horizon Discovery (for human PCLS). All JAK inhibitors were purchased from SelleckChem. qPCR primers were purchased from Integrated DNA Technologies. All human sample research was approved by the Brigham and Women’s Hospital Institutional Review Board.

### Computational analysis

Bulk RNA-seq dataset of human IPF lungs (N=231) and control lungs (N=267) was obtained from phs001662 (Lung Tissue Research Consortium (26)) and plotted for the expression level of genes in the IL-6 family.

Single-cell RNA-seq dataset of human IPF lungs (N=35) was obtained from GSE136831(27) and analyzed via Seurat (v. 4.0.4) for percent of cells expressing a feature, such as LIF, log-normalized and scaled average expression of a feature in a cell population.

### Cell Stimulation and Antibody Blocking Assays

Fibroblasts were plated on day 1 at 10,000 cells per well in 96-well plates in 10% FBS containing media. Cells were serum-starved on day 2 by changing to 1% FBS-containing media. Cells were stimulated as indicated on day 3 or blocking antibodies were added 1 hour prior to cytokine stimulation on day 3.

### siRNA Silencing

Fibroblasts were transfected with an siRNA by reverse transfection at 30 nM using the RNAi Max reagent (Life Technologies) in 10% FBS containing media. Cells were then switched to serum-starving media containing 2% FBS on day 2. Cells were stimulated as indicated on day 3. For human PCLS, siRNA was added to the media on day 0. On day 2 and 4, PCLS were snap-frozen in liquid nitrogen before RNA samples were extracted.

### Quantitative Real-Time PCR

mRNA collection and cDNA synthesis from fibroblasts was carried out using the Power SYBR Green Cells-to-Ct Kit (Life Technologies). mRNA samples were extracted from grounded PCLS using the RNeasy Fibrous Tissue Mini Kit (Qiagen) and cDNA synthesis was carried out using the QuantiTect Reverse Transcription Kit (Qiagen). qPCR reactions were performed using the Brilliant III Ultra-Fast SYBR reagent (Agilent). Relative transcription level was calculated by using the Δ ΔCt method with GAPDH (for fibroblast cells) and HPRT (for PCLS) as the normalization control.

### Western Blotting

Total cell lysates were collected by washing cells once with cold PBS followed by addition of lysate buffer (50mM HEPES pH 7.5, 5% glycerol, 100 mM NaCl, 0.1% SDS, 1% Triton X-100, supplemented with protease inhibitors and phosphatase inhibitors sodium orthovanadate, sodium fluoride, and beta-glycerol phosphate). Cells were lysed for 30 minutes on ice followed by centrifugation at 14000 rpm for 15 minutes at 4oC. Protein concentration was measured by the microBCA kit (Pierce). Equal amounts of total protein (∼20 µg per lane) were separated on an 8% SDS-PAGE gel. Proteins were transferred onto a PVDF membrane and blocked with 5% BSA in PBS and probed with primary antibodies overnight at 4oC, followed by secondary antibodies conjugated with HRP. Membranes were developed with the Clarity Western ECL Substrate (Bio-rad) and scanned with the ChemiDoc Imaging System (Bio-rad).

### Whole mount imaging

PCLS were washed with PBS-T (0.2% Triton X-100 in 1x PBS) and blocked for 1hr using 5% Goat-serum (GS) containing PBS-T with 3% BSA. PCLS were then incubated overnight at 4ºC with primary antibody solution (5% GS, 3% BSA in PBS-T). Next, PCLS were washed and incubated overnight at 4ºC with a secondary antibody solution (5% GS, 3% BSA in PBS-T). PCLS were then cleared with BABB (Benzyl Alcohol/ Benzyl Benzoate) before imaging.

## Author Contributions

This study was conceived and designed by HNN, YJ, EYK, and MBB. The study was supervised by EYK and MBB. The bioinformatic analysis was performed by YJ and supervised by EYK. The bioinformatic analysis of LTRC samples was performed by YJ in collaboration with JHY, PC, CPH. The clinical biobanking was performed by HA, YJ, Yaunghyun Kim, LMS, MV and supervised by RFP, LMS, MV, NSS, and EYK. The in vitro experiments were performed by HN, YJ, and Yaunghyun Kim. The PCLS immunohistology was performed by Yunhye Kim and supervised by KY. The manuscript was written by HNN, LTM, EYK, MBB and was revised by HNN, YJ, Yunhye Kim, Yaunghyun Kim, HA, LMS, MV, NSS, JHY, LTM, KY, EYK, MBB. Project administration by EYK and MBB. All authors have approved the submitted version.

## Classification

Biological Science - Immunology and Inflammation

## Acknowledgments

We thank the assistance of the BWH Tissue Bank. We also thank Dr. Jean-Francois Moreau and Dr. Vincent Pitard for providing valuable reagents. This work was supported by the BWH Bell Family Fund (EYK), American Lung Association Dalsemer Award DA-827785 (EYK), National Institute of Health (NIH) R01AR073833, P01AI148102 AND R01AR06370 (MBB). Molecular data for the Trans-Omics in Precision Medicine (TOPMed) program was supported by the National Heart, Lung and Blood Institute (NHLBI). RNASeq for “NHLBI TOPMed: LTRC” (phs#001662) was performed at Northwest Genomics Center (HHSN268291600032I). Core support including centralized genomic read mapping and genotype calling, along with variant quality metrics and filtering were provided by the TOPMed Informatics Research Center (3R01HL-117626-02S1; contract HHSN268201800002I). Core support including phenotype harmonization, data management, sample-identity QC, and general program coordination were provided by the TOPMed Data Coordinating Center (R01HL-120393; U01HL-120393; contract HHSN268201800001I). We gratefully acknowledge the studies and participants who provided biological samples and data for TOPMed.

