## Supplemental data for "Leukemia inhibitory factor (LIF) receptor amplifies pathogenic activation of fibroblasts in lung fibrosis"

* Edy Y Kim

* Michael B Brenner

**This PDF file includes:**

Figures and legends for S1 to S7

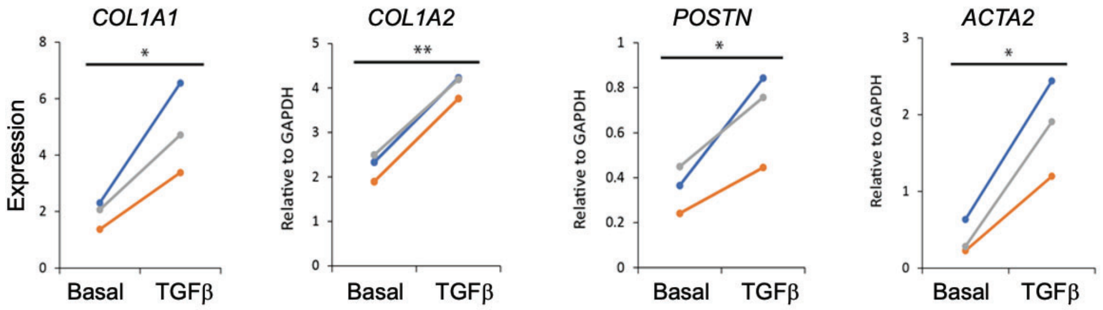


**Fig. S1.** TGFβ1 induces profibrotic genes in IPF lung fibroblasts. Early passage, primary lung fibroblasts were derived from IPF patients. Each line represents a different patient. Data represent gene expression in fibroblasts at 24 h after stimulation by TGFβ1 as measured by qPCR and normalized to GAPDH. (A) Fibroblasts were incubated with TGFβ1 (5ng/ml) for 24 h. Paired Student's t test. *p < 0.05, **p < 0.01.

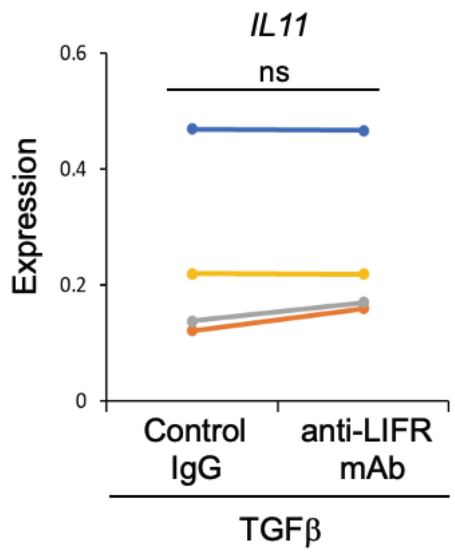


**Fig. S2.** Blockade of LIFR in lung fibroblasts did not suppress the TGFβ-dependent induction of *IL11*. Primary IPF lung fibroblasts were stimulated with TGFβ1 (5ng/ml) in the presence of an antibody against LIFR (LIFR mAb) or an isotype control (Ctrl IgG). Data represent gene expression in fibroblasts at 24 h after stimulation by TGFβ1 as measured by qPCR and normalized to *GAPDH*. Each line represents a different patient. Paired Student's t test. *$p<0.05$, ns: non-significant.

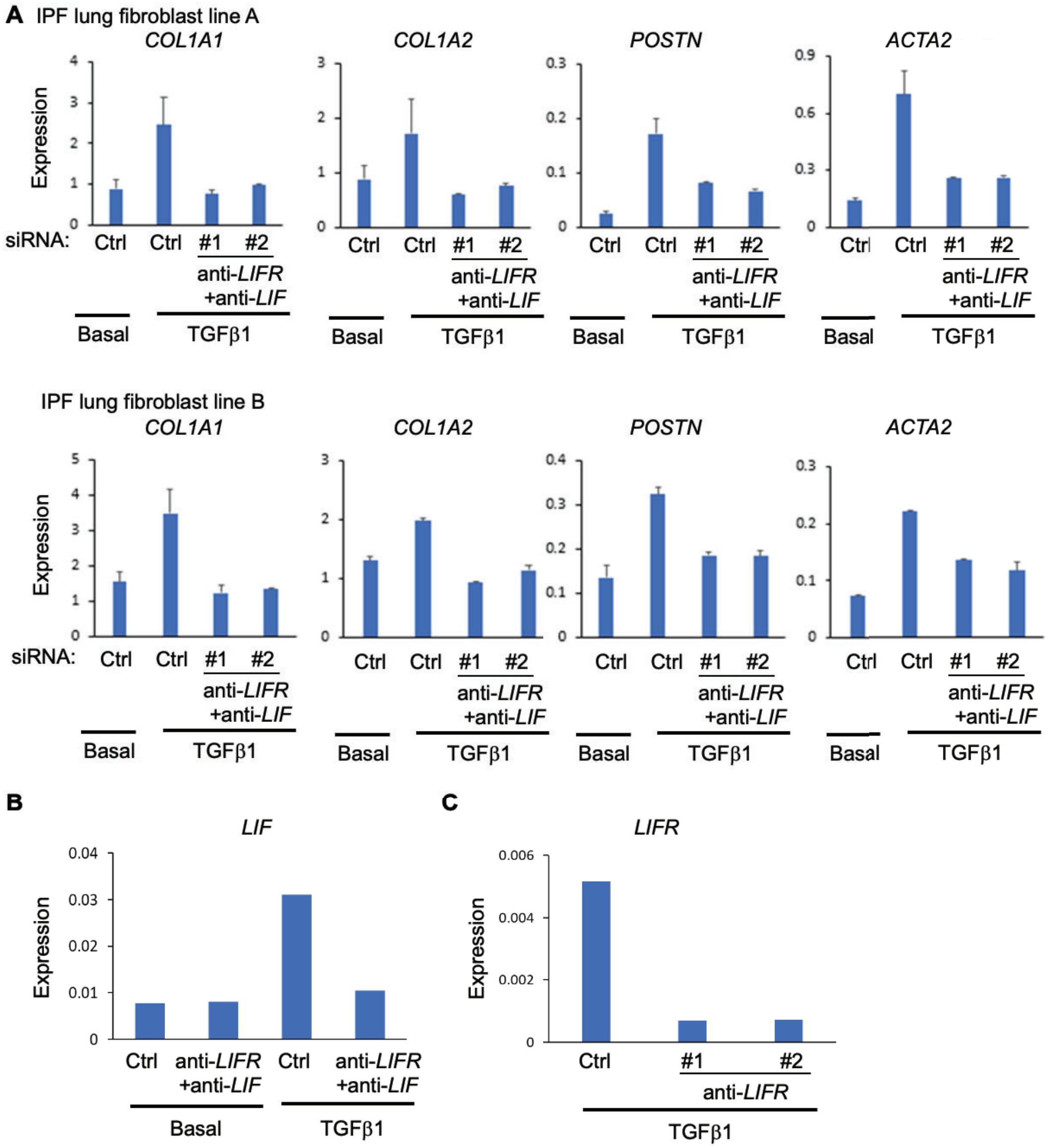


**Fig. S3**. Blockade of the LIF / LIFR autocrine loop with siRNA prevented the expression of TGFβ1-induced profibrotic genes. Primary IPF lung fibroblasts were transfected with control siRNA (Ctrl) or with siRNA against *LIF* and one of two different siRNA against *LIFR* (LIFR #1 or #2). After 48 h after siRNA transfection, fibroblasts remained in media (basal) or were stimulated with TGFβ1 (5ng/ml). 24 h after stimulation with TGFβ1, gene expression (normalized to *GAPDH*) were measured by qPCR. (A) Expression of profibrotic genes in lung fibroblasts from two IPF patients (lines A and B). (B) Expression of *LIF*. (C) Expression of *LIFR*. ANOVA with post-hoc testing. *p<0.05, ns: non-significant.

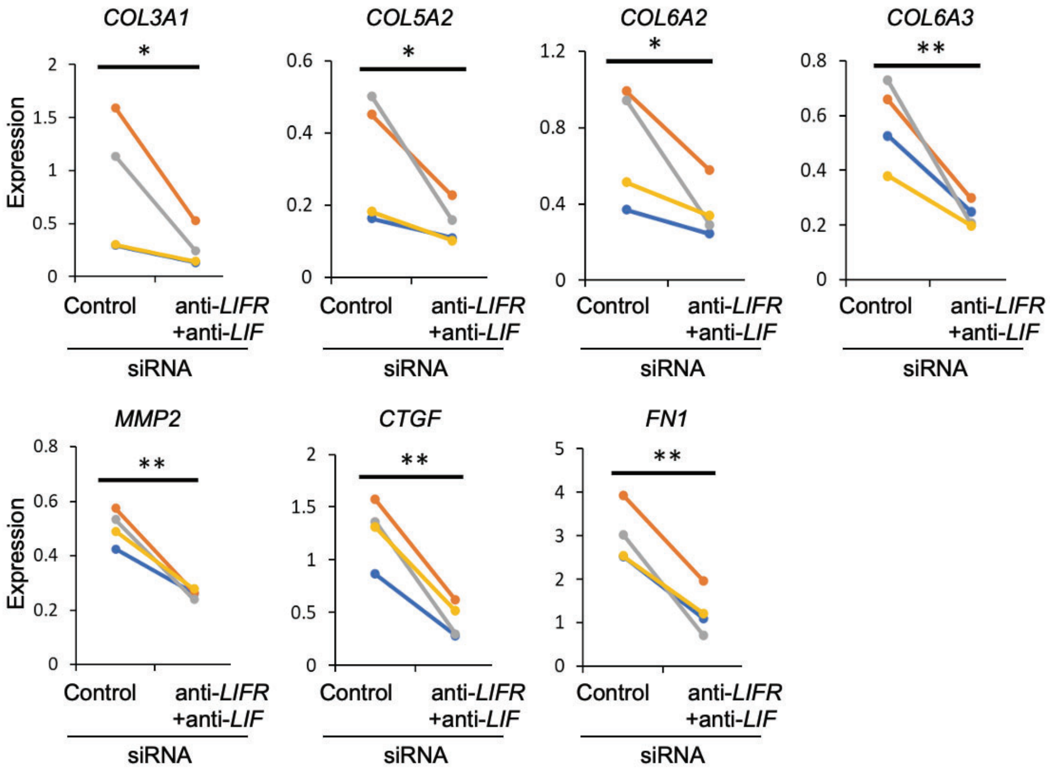


**Fig. S4.** Blockade of the LIF / LIFR autocrine loop with siRNA prevented the expression of additional TGFβ1-induced profibrotic genes. Primary IPF lung fibroblasts were transfected with control siRNA (Ctrl) or with siRNA against LIF and LIFR (LIFR #1). After 48 h after siRNA transfection, fibroblasts were stimulated with TGFβ1 (5ng/ml). 24 h after stimulation with TGFβ1, gene expression (normalized to GAPDH) were measured by qPCR. Each line represents a different IPF patient. Paired Student's t test, *p < 0.05, **p < 0.01, ***p < 0.001.

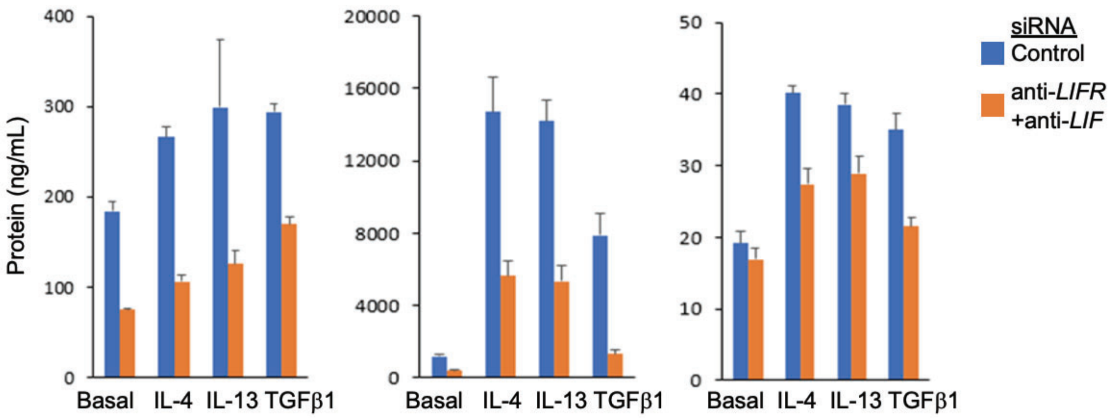


**Fig. S5. LIFR signaling regulates profibrotic gene programs**. Primary IPF lung fibroblasts were transfected with siRNAs against *LIFR* (siRNA LIFR #2) and *LIF* (LIFR + LIF) or a control (Ctrl) siRNA. 48 h after transfection with siRNA, cells were stimulated with IL-4 (50ng/mL), IL-13 (50ng/mL), TGFβ1 (5ng/ml), or left unstimulated (basal). The protein levels were measured by ELISA in cell lysates at 72 h post-stimulation. Error bars represent SD of quadruplicate technical replicates.

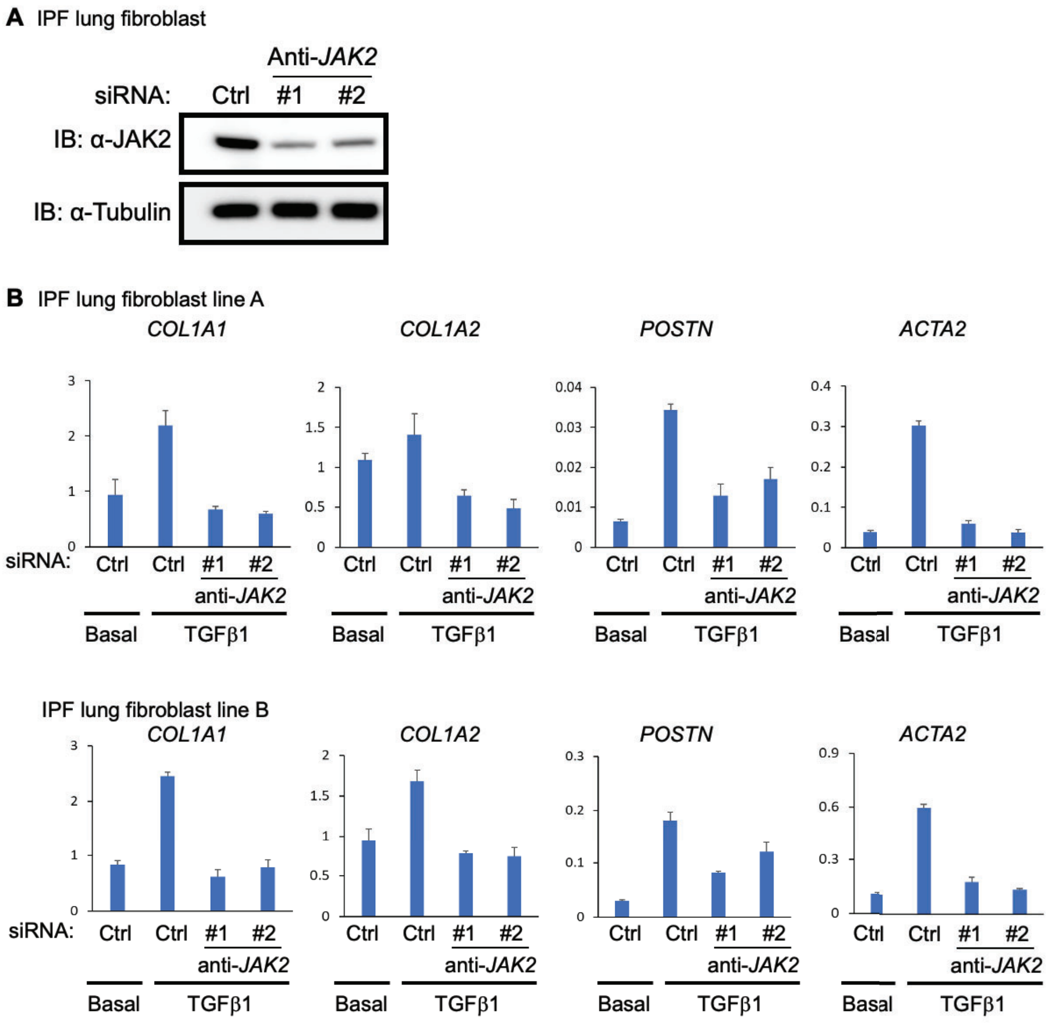


**Fig. S6.** JAK2 is required for induction of profibrotic genes by TGFβ1 in IPF lung fibroblasts. Primary IPF lung fibroblasts were transfected with control (Ctrl) siRNA or one of two different siRNA against JAK2 (JAK2, #1 or #2). 48 h after siRNA transfection, fibroblasts were stimulated with TGFβ1 (5ng/ml). 24 h post-TGFβ1 stimulation, gene expression (normalized to GAPDH) were measured by qPCR. JAK2, #1 siRNA was used in Figure 4C.

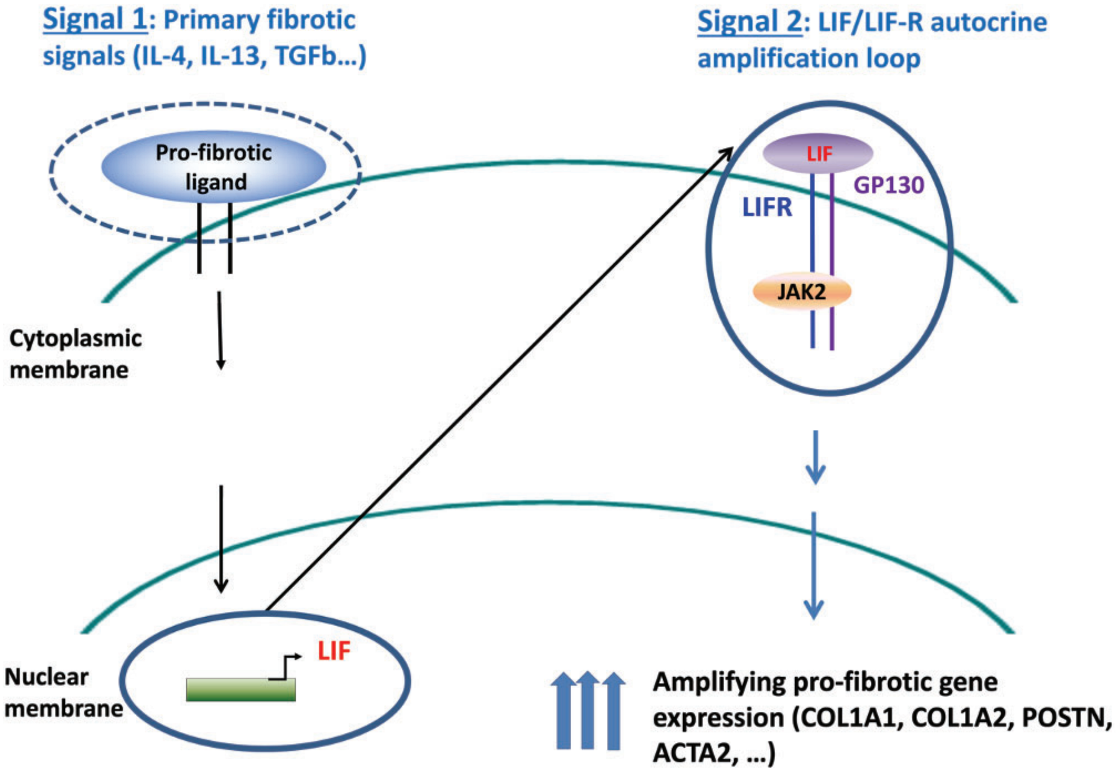


**Fig. S7.** Schematic of autocrine LIF/LIFR as a master amplifier of fibroblast mediated fibrotic gene expression. In this schematic, after a range of profibrotic stimuli ("Signal 1"), fibroblasts upregulate LIF. Next, in an autocrine loop, LIF ligand binds to LIFR on the same fibroblast ("Signal 2"), which then drives and amplifies the profibrotic gene program (blue arrows).
